# Condition-dependent trade-offs maintain honest signaling: A laboratory experiment

**DOI:** 10.1101/788828

**Authors:** Szabolcs Számadó, Flóra Samu, Károly Takács

**Affiliations:** Department of Sociology and Communication, Budapest University of Technology and Economics, Egry J. u. 1. H-1111, Budapest, Hungary; CSS-RECENS, Centre for Social Sciences, Hungarian Academy of Sciences, Tóth K.u. 4, H-1097, Budapest, Hungary; Evolutionary Systems Research Group, MTA, Centre for Ecological Research, Hungarian Academy of Sciences, Klebelsberg Kunó str. 3., 8237 Tihany, Hungary; The Institute of Analytical Sociology, Linköping University, Norra Grytsgatan 10, S-601 74 Norrköping, Sweden; Corvinus University of Budapest, Doctoral School of Sociology

**Keywords:** honest signaling, Handicap Principle, costly signaling, signaling trade-offs, condition dependent trade-offs

## Abstract

How and why animals and humans signal reliably is a key issue in biology and social sciences. For many years the dominant paradigm in biology was the Handicap Principle. It claims a causal relationship between honesty and signal cost and thus predicts that honest signals have to be costly to produce. However, contrary to the Handicap Principle, game theoretical models predict that honest signaling is maintained by condition dependent signaling trade-offs and honest signals need not be costly at the equilibrium. Due to the difficulties of manipulating signal cost and signal trade-offs there is surprisingly little evidence to test these predictions either from biology or from social sciences. Here we conduct a human laboratory experiment with a two-factorial design to test the role of equilibrium signal cost vs. signalling trade-offs in the maintenance of honest communication. We have found that the trade-off condition has much higher influence on the reliability of communication than the equilibrium cost condition. The highest level of honesty was observed in the condition dependent trade-off condition as predicted by recent models. Negative production cost, i.e. fix benefit-contrary to the prediction of the Handicap Principle-promoted even higher level of honesty than the other type of costs under this condition.

## 1. Introduction

Honest signaling under a conflict of interest is challenging to explain. The Handicap Principle dominated the biological literature in the past 30 years (Zahavi, 1975; Grafen, 1990; Zahavi & Zahavi, 1997). It claims that signals are honest because they are costly to produce (Zahavi, 1975; Grafen, 1990; Zahavi & Zahavi, 1997). While the Handicap Principle remains ill-supported (yet still dominant) in biology (John, 1975; Borgia, 1979; Kotaiho, 2011; Getty, 2006; Számadó, 2011; Groose, 2011; Számadó & Penn, 2015, 2018; Penn & Számadó, 2019) many found applications of it in explaining human behaviour from Potlach to big-game and turtle hunting (). However, game theoretical models predict that it is not the equilibrium cost that maintains the honesty of communication but the ratio of marginal costs to marginal benefits (Spence, 1973; Riley, 1979; Hurd, 1995; Számadó, 1999; Lachmann et al., 2001; Bergstrom et al., 2002). This means that honest signals need not be costly to produce at the equilibrium not even under conflict of interest (Számadó, 1999; Lachmann et al., 2001; Számadó, 2011; Higham, 2014), cost-free or even negative cost signals can be honest and evolutionarily stable (Enquist, 1985; Hurd, 1995; Számadó, 1999 Lachmann et al., 2001; Számadó, 2011; Számadó et al., 2019). In line with this current trend anthropological reviews challenge the dominance of Handicap Principle and emphasise that signals used by humans need not be overly costly to be honest and useful (B, Stibbard). All in all, there seems to be an unresolved issue whether signals used by humans need to be costly to produce to be honest, as predicted by the Handicap Principle, or not, as predicted by models and recent reviews of the anthropological literature. Here we test these two conflicting proposals in a laboratory experiment where participants play a simple signaling game. The experiment has a two-factorial design, we varied both the absence or presence of signal trade-offs and the equilibrium cost of signals for honest signalers including negative, zero and positive equilibrium cost. If the Handicap Principle is correct, signals will be honest only in the positive equilibrium cost condition. If game theoretical models are correct then signals will be honest only in the condition-dependent trade-off condition. Our results support the latter claim: honest signaling emerged under the condition-dependent trade-off condition regardless of the equilibrium cost of signals; i.e. signals with zero or even negative production cost (benefit) were honest under this condition. Our results demonstrate that the conditions that allow the emergence of honest signalling is different from and much wider than what the Handicap Principle would dictate.

## 2. Methods

We used a two-factorial design where we varied both the presence of signaling trade-offs and the value of equilibrium cost for honest signalers (see Table 1). In the trade-off conditions, we test whether the absence, the presence or differentiation between the costs of the two signals regarding the types of signalers influence the emergence of honest equilibrium. We had three trade-off conditions: (i) no trade-off, where the cost of low vs. high signal was the same regardless of the condition of signaler; (ii) trade-off condition, where sending the high signal was costlier for both low and high condition signalers in the same way (i.e. the marginal cost of sending a high signal was the same for all conditions); (iii) condition-dependent trade-off condition, where the cost of sending a high signal was costlier for signalers in low condition (i.e. the marginal cost of sending a high signal was higher for signalers in low condition, i.e. Grafen’s (1990) equations (b) and (c) p.521). In order to prove the effects of different trade-offs on the emergence of honest communication, regardless of whether the honest signals are costly or beneficial for the high condition signalers, we test the trade-off manipulations within three equilibrium cost condition: (i) positive equilibrium cost, where high condition signalers had to pay to use the high signal; (ii) zero equilibrium cost where high condition signalers could use the high signal for free; (iii) negative cost condition, where high condition signalers received a benefit (payment) for using the high signal.

**Table 1.** Treatment conditions. The table shows the costs and benefits signalers receive for using low intensity or high-intensity signals. Costs and benefits depend on the signalers’ condition. In this table, positive numbers are costs, negative numbers indicate benefits. First of all, the treatments differed in whether between the two signal has no contrast (trade-off = 0) or switching between the two signals costs the same (trade-off = 1500), and in the third case, it is lower for the signallers in high condition (trade-off=200). Second, treatments also differed in whether the use of the high-intensity signal is costly, cost-free or even profitable for the high condition signallers. (c_11_ for low-intensity signal, c_12_ for high-intensity signal) (c_21_ for low-intensity signal, c_22_ for high-intensity signal)

|  | equilibrium cost | trade-off | cost for signalers in low condition | cost for signalers in high condition | Expectation |
| --- | --- | --- | --- | --- | --- |
| 1. | negative | no | (0, 0) | (-100, -100) | dishonest |
| 2. | negative | yes | (0, 1500) | (-1600, -100) | dishonest |
| 3. | negative | condition d. | (0, 1500) | (-300, -100) | honest |
| 4. | zero | no | (0, 0) | (0, 0) | dishonest |
| 5. | zero | yes | (0, 1500) | (-1500, 0) | dishonest |
| 6. | zero | condition d. | (0, 1500) | (-200, 0) | honest |
| 7. | positive | no | (0, 0) | (100, 100) | dishonest |
| 8. | positive | yes | (0, 1500) | (-1400, 100) | dishonest |
| 9. | positive | condition d. | (0, 1500) | (-100, 100) | honest |

One experimental session contains three treatment conditions, and in each condition, participants played 20 rounds of the signaling game. In the repeatedly played rounds participants alternately act as a signaler or as a receiver. Because of the alternating roles, players were informed about the cost of each signal right before the game (see Further Instructions in Appendix II). With full knowledge of the signal costs the expected equilibria (see hypotheses below) became clear to the players. This is not problematic, because in this experiment we are less interested in whether participants are able to find the separating equilibrium, but our interest rather relates to its maintenance: the question here is indeed, under what conditions the separating equilibrium can be maintained in a signaling game with conflict of interests, where different players have different equilibrium preferences: R and S in high condition have higher payoffs in separating equilibrium, in contrast, S in low condition has higher outcome on average in the pooling equilibrium or an unbalanced state.

### Predictions

In a condition where the cost of the two signal is equal for both types of signalers – or in other words where there is no trade-off between the signals – an equilibrium cannot evolve since there are no such individual strategies from which it is not beneficial for either party to move towards another action.

*H1. In the absence of trade-off, the signalers are expected to select randomly from the two available signals, therefore receivers respond also randomly to the signals they receive.*

Costs of the signals in the second condition – where there is an equal trade-off between the two signal – are different. They are configured in a way that, on the one hand, for S in low condition the cost of using the high-intensity signal (c_12_) exceeds the benefit of the resource from R (1500>b, b=1200), on the other hand – since the trade-off between the two signals is equal for both S – for S in high condition the benefit of using the low-intensity signal (c_21_) exceeds the benefit from R’s resource (1600>1500>1400>b, b=1200). The consequence of this we expect a pooling equilibrium, in which both signalers send the same signal (low-intensity signal) and therefore, R is unable to predict the type of the signaler.

*H2. If both types of signalers have the same trade-off between the two signals, they will use the same signal, and receivers decide randomly on the allocation of the source.*

If the trade-off between the two signals is condition dependent – meaning its marginal cost is different for the different types of players S – S in high condition will switch to the “high intensity” signal in order to acquire resources from R since in this case the marginal cost of moving to this signal is smaller than the marginal benefit of it.

*H3. If the two signalers have a different trade-off between the two signals they will use different signals depending on their type, thus the receivers will be able to determine the type of signalers correctly and allocate the resources according to their interest.*

## 3. Results

We go through the results in the same order as we formulated the hypotheses. We look at the aggregate results first, and then we check their validity with more complex methods. In this section three outputs will be examined: (1) to what extent are signals with different intensity separated by the signalers’ condition, (2) what are the probabilities of resource allocation according to which signal was seen, and (3) the overall degree of coordination between response and condition, i.e. to what extent has the condition of the signalers been successfully determined by the receivers.

H1. In the absence of trade-off, the signalers are expected to select randomly from the two available signals, therefore receivers respond also randomly to the signals they receive.

Figure 2 shows that different types of signalers used the two signals with the same probability in treatments where there was no difference in the cost of using the two signals (no trade-off), regardless of whether the signals were positive, negative or no cost (see graphs in the first column in Figure 2). Looking at the receiver’s responses to these random signals, we also see that there was no difference in these treatments in the extent to which the receiver gave the resource after seeing one or the other signal. (see graphs in the first column in Figure 3). Finally, in these treatments, half of the time (49%) R estimated signalers’ original states correctly (gave the resources to S in high condition and refuse to give it to S in low condition) (see Figure 4).

**Figure 1.**
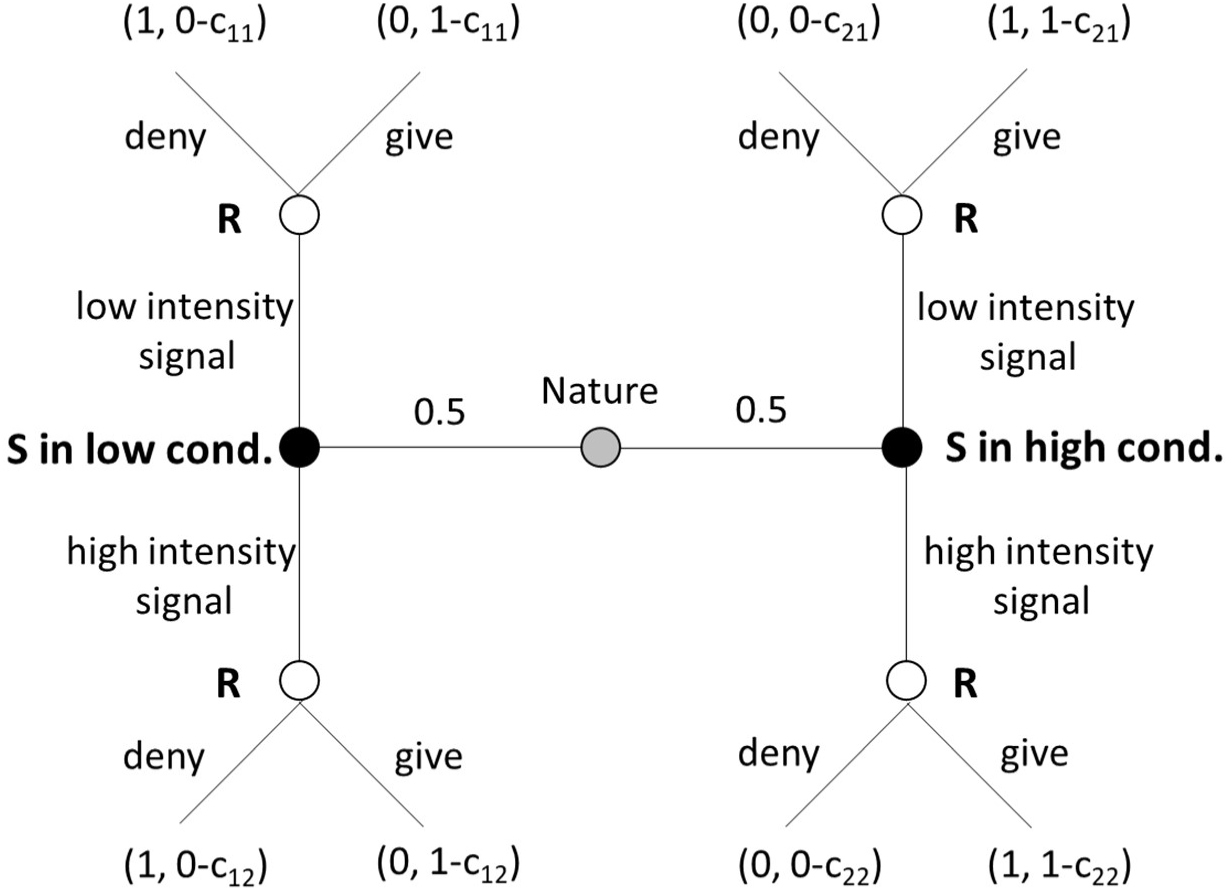
The signaling game with conflict of interest, differential cost model (i.e. the cost is condition dependent) After nature randomly divides the roles (S and R) and the types of signalers (S in high and in low condition) players make their decisions successively. First, S can send one from two different signals (low-intensity signal, high-intensity signal) to get the resource from R. After this, R gets the signal and decides whether to provide the resource to a given signal or not. At the end of each round players get feedback on their success. S both in high and low condition win when they get the resource from R, but for R only the dedication of the resource to S in high condition and its protection from S in low condition generate successful outcomes (see outcomes in brackets at the ends of the decision tree where the first element refers to R and the second to S). In the figure c_11_, c_12_, c_21_, and c_22_ indicate the different cost of signals according to the type of signaler. The figure also contains the codes used in the experiment.

**Figure 2.**
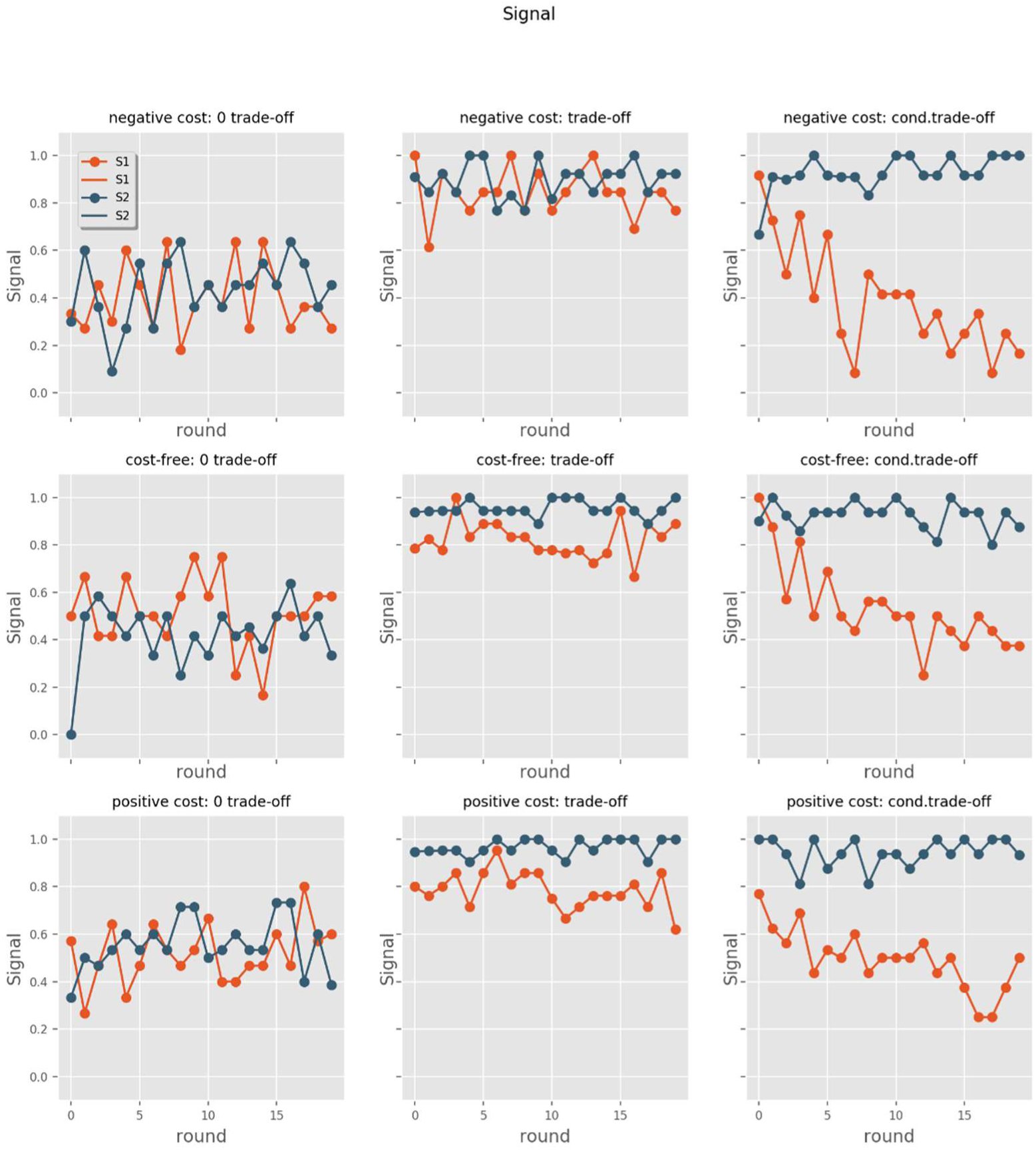
Signalers’ decisions by rounds, by the types of the signalers and by the nine treatments. The Y-axis shows the probability of a low-intensity signal has been sent. Sending high intensity is complementary to this. The lower the line, the higher the probability of a high-intensity signal has been sent. The blue line shows the decision of signalers in low condition, the orange is for signalers in high condition. The figure shows that there is a difference between columns rather than between rows, which means that trade-off conditions tend to influence the development of honest communication rather than the type of the equilibrium cost for high condition signalers for using high-intensity signals.

**Figure 3.:**
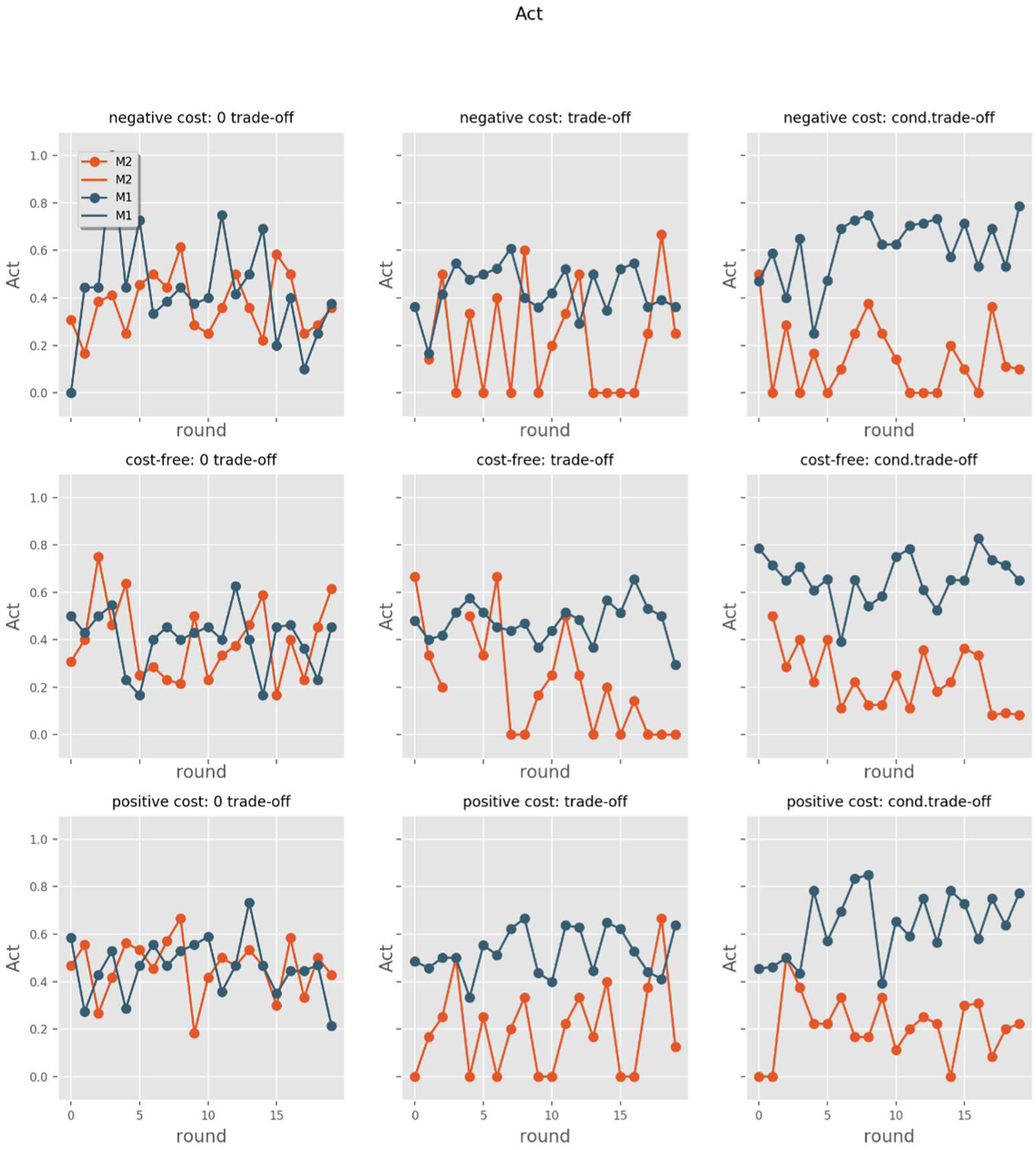
Receivers’ decisions by rounds, by the observed signals and by the nine treatments. The Y-axis shows the probability of the resource has been denied. Giving the resource is complementary to this. The lower the line, the higher the probability of giving the resource. In this figure, it can be also noticed that patterns differ along with the trade-off conditions, while the lines are quite similar to each other.

**Figure 4.**
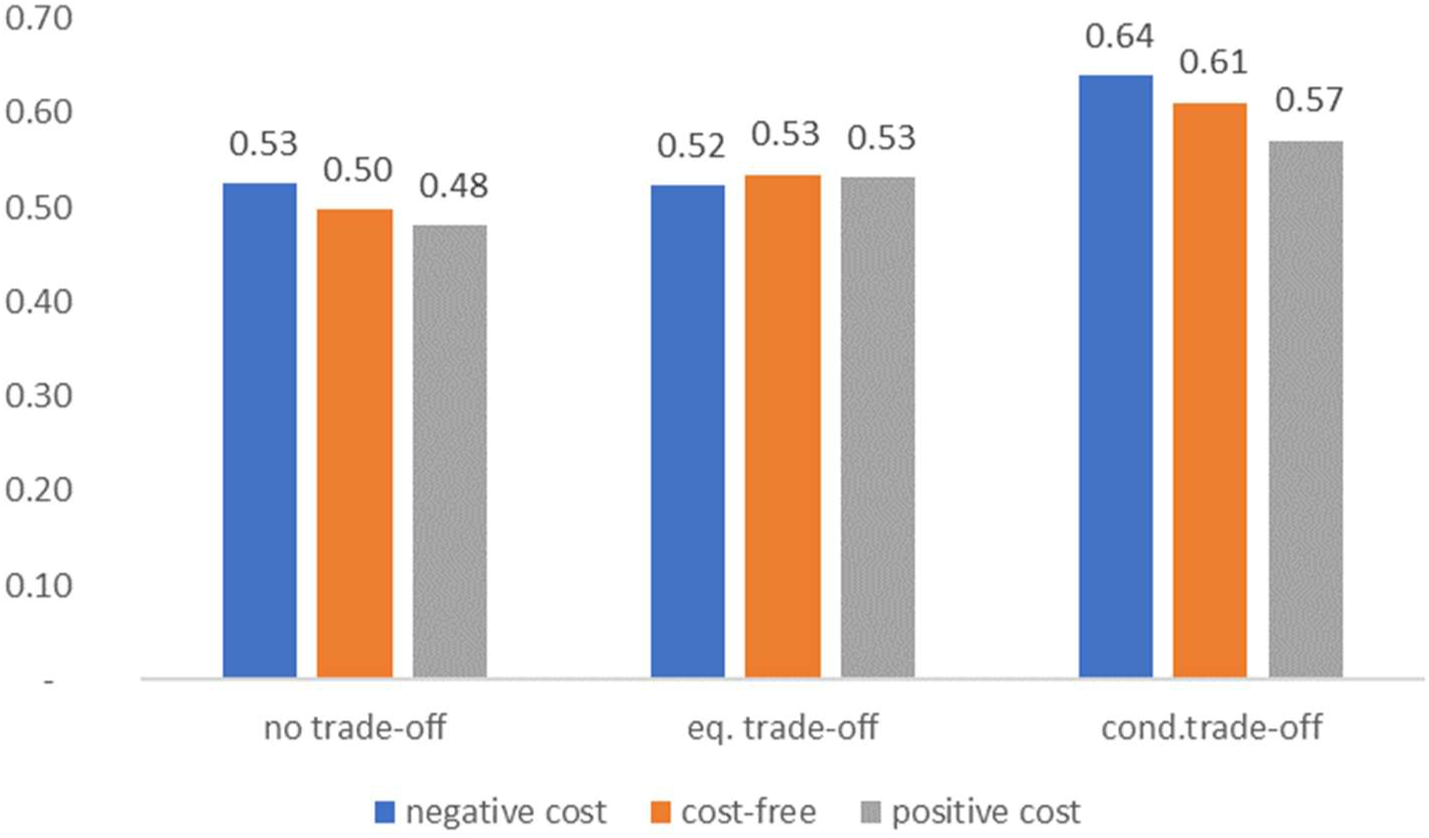
The overall success of Rs by treatments. Although there is no significant success rate, there is an increase in trade-off treatments, especially in the conditional-dependent trade-off treatments. Ratios are probably independent of the different equilibrium costs.

H2. If both types of signalers have the same trade-off between the two signals, they will use the same signal, and receivers decide randomly on the allocation of the source.

In treatments with the same trade-off for both signalers, we see that the majority (88%) of signalers are using one signal (see the second column in Figure 2). It should be noted that it is a strong but not a perfect pooling equilibrium. We also see divergence within receivers’ decision by signals in these treatments (see the second column in Figure 3). Receivers tend to favor high-intensity signals (giving 80.8% of the resource) compared to low-intensity signals (51.6%). Here R’s success ratio was slightly higher than in the previous treatment (53%). (see Figure 4)

H3. If the two signalers have a different trade-off between the two signals they will use different signals depending on their type, thus the receivers will be able to determine the type of signalers correctly and allocate the resources according to their interest.

Results observed in the third trade-off manipulation are also consistent with our expectations. Signals diverge with the greatest extent by the type of signaler in these treatments. Learning dynamic can be observed in the last column in Figure 2 where high condition signalers gradually switch to the use of a signal with favourable trade-off to them (which need not be costly) to achieve coordination. In this treatment, players move much more towards separating equilibrium, as marginal benefits for high condition signalers still outweigh the marginal cost of more expensive signals. Most of the players in this role used high-intensity signal in the last rounds of the game while more than 90% of the signalers in low condition were stuck to the low-intensity signal. On the other side, 80.4% of the resource was given if receivers saw the high-intensity signal and almost two-thirds of the low-intensity signals were rejected (63.8%) (see figures in the last column in Figure 3). In terms of the overall success, R’s decisions were the most successful in this treatment (60%) which is not yet perfect but definitely better than in previous treatments (see Figure 4).

Within the trade-off treatments, we see similar patterns among the manipulation of equilibrium cost (negative, cost-free, positive), therefore we think equilibrium cost has no effect on honest communication.

We run more complex models to test whether this shift towards a separating equilibrium can be detected statistically after we control for other variables besides the number of participants and the order of the games (see Regression tables in Appendix I). We used mixed-effects logistic regressions to deal with the problem of observing the same people through multiple rounds. The regression models revealed the same results as the aggregated figures. R decided most accurately in the condition-dependent trade-off treatment (see *cond. trade-off* effect in Table 1, Appendix I.), where signalers in high condition were more likely to transmit high intensity signals as time passed (round effect in Model 3, Table 2, Appendix I.), while conversely, low condition signalers’ tendency of sending low-intensity signals increased over time (see *S in low condition # round* interaction effect in Model 3, Table 2, Appendix I.). Although we can see a significant difference in signaling between the two types of signalers also in the simple trade-off treatment, this difference does not intensify over time, which means, it is not directed towards separating equilibrium, as opposed to the condition-dependent trade-off treatment (see round and interaction effects in Model 2, Table 2, Appendix I.). Negative equilibrium cost facilitates sending high-intensity signals compared to positive equilibrium cost, meaning participants switch more easily between signals if they still win a small amount of money than if they lose by using high-intensity signals.

We can observe order effect in the no-trade off treatments, where high-intensity signal was more likely has been sent when the game was played after a game with honest equilibrium (see Model 1, Table 2, Appendix I.). In the consecutive games, different signal pairs were used, the order effects can be interpreted as participants might send messages less randomly. Playing additional games intensify the use of low-intensity signals in the no trade-off treatment, and the high-intensity signals in the simple trade-off treatment where costs helped players to find out which signal is different from the one that leads to a pooling equilibrium. As the number of players increases, the shift to the separating equilibrium slows down in the condition-dependent trade-off condition, and to the pooling equilibrium in the simple trade-off treatment. (see the effect of N in Model 2 and 3, Table 2, Appendix II.)

When it comes to resource allocation for low-intensity signal less resource was given in both trade-off treatments, but this separation grew over time only in the condition dependent trade-off treatment. (see low-intensity signal # round interaction effect in Model 2 and 3, Table 3, Appendix II.)

This divergence may stem from random errors, but the pattern suggests participants may calculate that part of their final payoffs will be selected from those rounds where they acted as R, and therefore they tried to move toward the separating equilibrium by choosing the costlier (less beneficial) signal when they are acting as high condition signaler since it leads to the separating equilibrium where R can make more precise decision.

This failing rate can be explained by accidental errors or the relative shortness of the game.

## 4. Discussion

Honest signaling emerged in our experiment under the condition-dependent trade-off condition regardless of the equilibrium cost of signals. The highest level of honesty was achieved under the condition-dependent trade-off condition with negative signal cost. No honesty evolved without trade-offs and some level of honesty evolved with simple trade-offs under the conditions of cost-free and costly equilibrium cost signals. Signals with negative signal cost seems to be counter intuitive. No honesty evolved under the simple trade-off condition and it took the longest time to find honest signalling under the condition-dependent trade-off condition. However, once it was discovered that signals with negative signal cost can maintain honesty-contrary to the predictions of the Handicap Principle and costly signaling models-the highest level of honesty was achieved under this condition. All in all, our results support the prediction of recent models that honest signaling is maintained by condition-dependent trade-offs instead of signal cost paid at the equilibrium.

Our results also show that while Grafen’s equations are technically correct (condition-dependent trade-off condition), his interpretation of these equations and his verbatim support for Zahavi’s Handicap Principle is incorrect (see more details Penn & Számadó, in press) since honesty emerged with zero or negative production cost. Honest signals need not be handicaps, in fact, the Handicap Principle is highly misleading in emphasizing the role of equilibrium cost instead of the role of signalling trade-offs (Számadó, 2011; Penn & Számadó, in press). Also, these trade-offs can be implanted between benefit function (Számadó & Penn,), i.e. having a cost function per se is not a requirement of honest signaling. The Handicap Principle is misleading both in its theoretical and empirical predictions, the sooner it is replaced with a Darwinian theory of honest signaling based on signaling trade-offs (Getty, 2006; Penn & Számadó, in press) the better.

## Methods

### Signaling game and honest equilibrium

In the 2×2 signaling game, there are two roles: the sender (S) and the receiver (R). R has the resources that S would like to receive. Two types of senders can be identified: S1 and S2. To give a better understanding we will call them hereafter S in *high* and in *low* condition. The type of the sender is important here since the signaling game what we used was based on the conflict between the receiver (R) and one of the signalers (S in low condition): R gets the highest payoff if it gives the resource to an S in high instead of S in low condition, however, both signalers are interested in receiving the resource from R (see Figure 1). Since R does not have information on which type of S sends the signal, R is trying to predict the type of the signaler for favorable resource allocation while S in low condition is trying to conceal it, because otherwise, with ‘honest’ communication, they would not receive the resource from R. Players can use different signals (call them hereafter *low* and *high* intensity signal) to gain the resource from R. By honest communication we mean the *separating equilibrium* of the signaling game when the two types of signalers use different signals which led to a consequence that signals take on the meaning of the type of the signaler (S in high condition or S in low condition). No meaning will evolve if signals do not differ by the type of the signalers, e.g. if both signalers send the same signal (*pooling equilibrium*). According to theoretical speculations (Spence 1973, Getty, 2005; Számadó, 2010; Számadó & Penn,; Számadó et al.,) the receiver can distinguish the type of signalers if the costs of the signals correlate negatively with the productivity of the signalers.

### Experimental setup

Participants played a simple 2×2 signaling game (see the description of the game in Figure 1). in a computer lab. 12 sessions were organized, involving different numbers of participants (groups of 12, 16, and 20). A total of 196 students participated in the experiment. A mix of within and between subject design was applied: within a session, each group played three of the nine treatment conditions. Each condition was played as the first, second and third game through 20 rounds each. Since the order of games and the number of participants (Bruner et al. 2014) may affect the speed of learning dynamics in the game, we control for these factors with statistical methods during the analysis. The experiment took place at the Corvinus University of Budapest (CUB) in Hungary between 25^th^ January 2018 and 15^th^ January 2019. Participants were regular or corresponding students and one experiment lasted for approx. 45 minutes. During the experiment, participants were seated randomly in front of the computers, thus participants took part in the experiment anonymously. Computers were connected on a local network with the help of the software z-Tree (Fischbacher 2007). The description of the game was included in the experimental instructions (see Appendix II.) and displayed on participants’ screens. They got the instructions on paper as well because roles participants played in the game have changed round-by-round. In the first step, participants were divided into groups of fours which contained two signalers (S) and two receivers (R). We used unbiased signaling game, where the two types of signaler (S in high condition and S in low condition) were assigned randomly by the computer. Everyone was only aware of their own type, they did not know each other’s condition. In the next step, signalers had to choose a signal and send it to the receiver to get the resource. Distinguishable characters were used as signals (e.g.)(and ∼) and their costs (or benefits) were displayed next to each signal depending on which condition was played. In the next step, receivers had to decide whether they would give the resource after a signal was seen. Signalers succeed if they receive the resource, but receivers’ success depended on the (hidden) type of the signaler (S in high condition was preferred to S in low condition). After these steps, participants have learned the type of the sender, the signal they sent and the success of their decision. In case of a successful decision, they received HUF 1200. The other factor that influenced the payments was the cost of the signals (see Table 1). In addition, participants received a show-up fee of HUF 1000.

## Acknowledgements

S.S., F.S. and K.T. was supported by the National Research, Development and Innovation Office – NKFIH (OTKA) grant K 112929, and by the European Research Council (ERC) under the European Union’s Horizon 2020 research and innovation programme (grant agreement No 648693). S.S. acknowledges support from by the National Research, Development and Innovation Office under grant number GINOP-2.3.2-15-2016-00057. The funding agencies had no role in the study design, analyses, or publication.

## Author contribution

S.S. conceived the idea, F.S. carried out the experiment, S.S. and F.S. analysed the results, S.S., F.S and K.T. wrote the paper.

## Appendix I. Regression tables

**Table 1.**
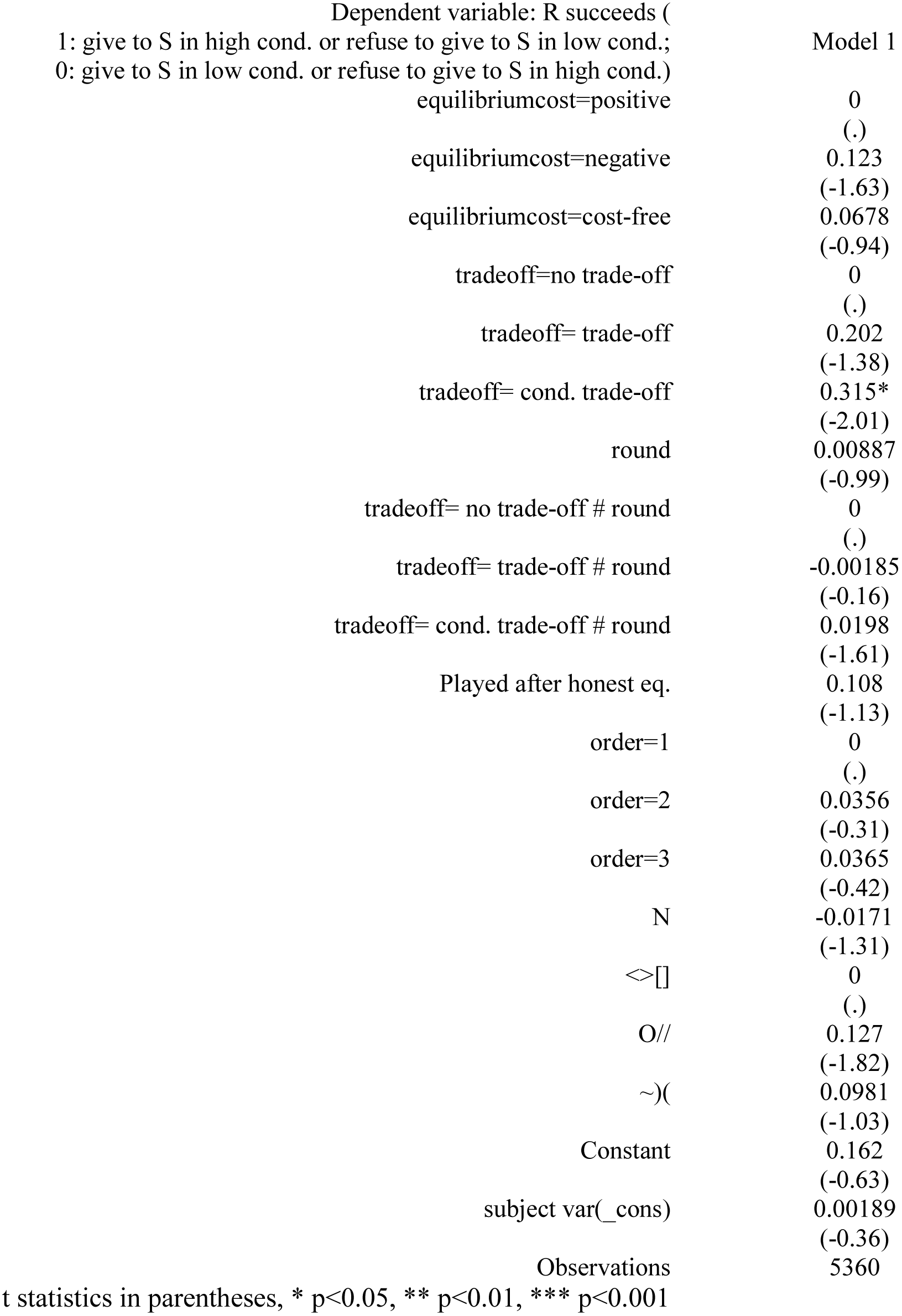
Mixed-effects logistic regression for R’s success

**Table 2.**
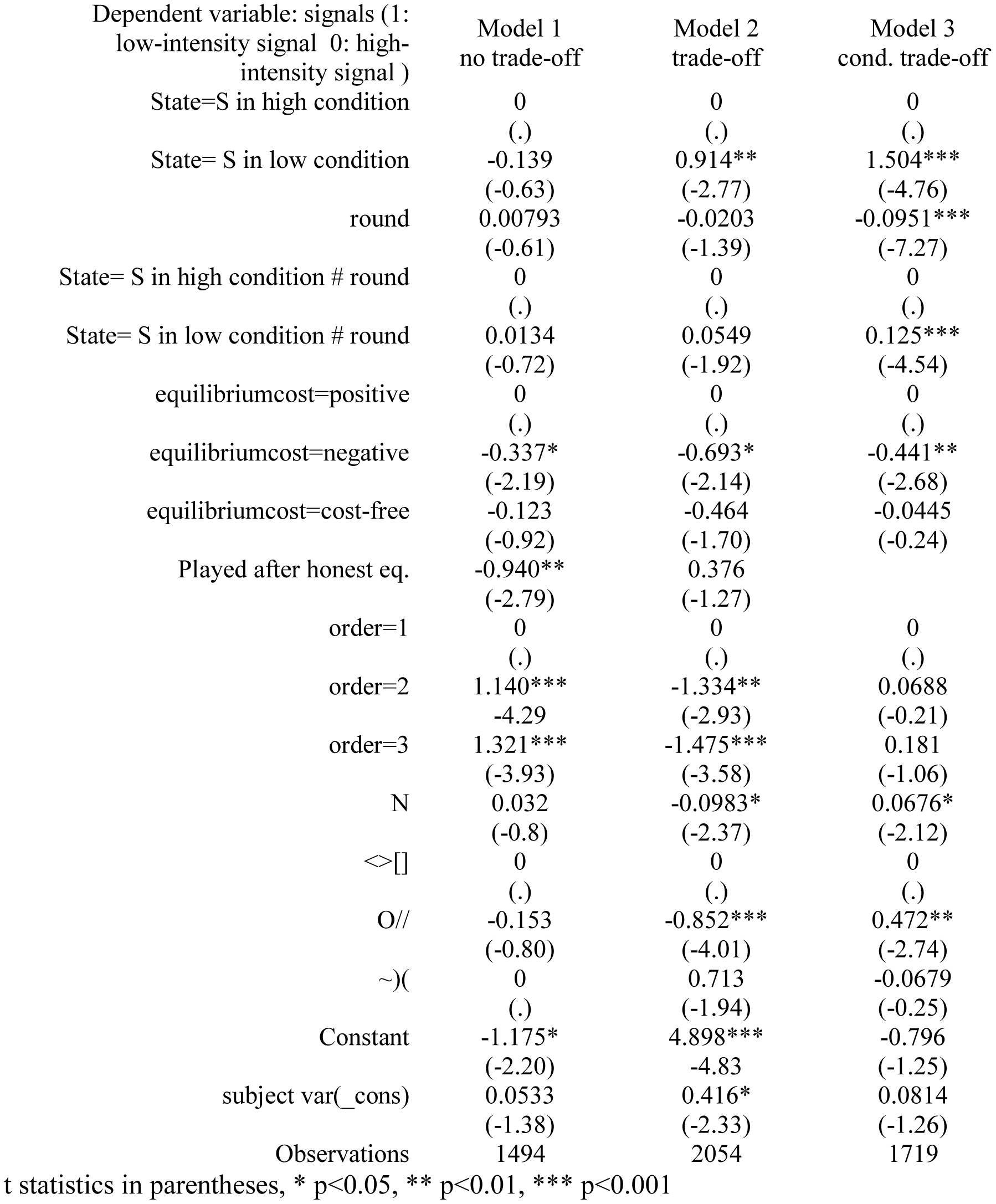
Mixed-effects logistic regression for S’s decision by trade-off manipulation

**Table 3.** Mixed-effects logistic regression for R’s decision by trade-off manipulation

| Dependent variable: R's<br>decision (1: refuse 0: give) | Model1<br>no trade-off | Model 2<br>trade-off | Model 3<br>cond. trade-off |
| --- | --- | --- | --- |
| Signal=high intensity signal | 0<br>(.) | 0<br>(.) | 0<br>(.) |
| Signal=low intensity signal | 0.219<br>(-0.96) | 1.082**<br>(-3.03) | 1.187***<br>(-4.11) |
| round | -0.00094<br>(-0.07) | -0.0215<br>(-0.76) | -0.0362<br>(-1.73) |
| Signal=high intensity signal #<br>round | 0<br>(.) | 0<br>(.) | 0<br>(.) |
| Signal=low intensity signal #<br>round | -0.0127<br>(-0.67) | 0.0298<br>(-1.01) | 0.0710**<br>(-3.01) |
| equilibriumcost=positive | 0<br>(.) | 0<br>(.) | 0<br>(.) |
| equilibriumcost=negative | -0.343*<br>(-2.21) | -0.00422<br>(-0.02) | -0.234<br>(-1.68) |
| equilibriumcost=cost-free | -0.304*<br>(-2.21) | -0.0127<br>(-0.09) | -0.0182<br>(-0.12) |
| Played after honest eq. | 0.3<br>(-0.89) | 0.322<br>(-1.75) |  |
| order=1 | 0<br>(.) | 0<br>(.) | 0<br>(.) |
| order=2 | -0.327<br>(-1.26) | 0.209<br>(-0.93) | -0.201<br>(-0.74) |
| order=3 | -0.449<br>(-1.33) | 0.227<br>(-1.06) | 0.165<br>(-1.18) |
| N | -0.0713<br>(-1.71) | -0.0308<br>(-1.18) | -0.00641<br>(-0.24) |
| <>[] | 0<br>(.) | 0<br>(.) | 0<br>(.) |
| O// | -0.065<br>(-0.33) | 0.0332<br>(-0.26) | 0.114<br>(-0.8) |
| ~)( | 0<br>(.) | -0.111<br>(-0.52) | 0.206<br>(-0.89) |
| Constant | 0.712<br>(-1.28) | -1.118<br>(-1.60) | -0.941<br>(-1.66) |
| subject var(_cons) | 0.154*<br>(-1.98) | 0.182*<br>(-2.3) | 0.015<br>(-0.65) |
| Observations | 1489 | 2041 | 1702 |

t statistics in parentheses, \* p<0.05, \*\* p<0.01, \*\*\* p<0.001

## Appendix II. Instructions

### INSTRUCTIONS

The experiment will consist of 60 rounds of decision-making, each round is one game. In each round, you will be paired with other participants. All participants are going to play the game simultaneously.

Before each round, the computer randomly pairs two players who will play together in that round. One member of the pair will be player X and the other will be player Y. In each round, these roles are also randomly assigned by the computer.

Moreover, there are two types of Y in the game: half of the Y players will be in a BLUE state and the other half will be YELLOW. These states are also randomly drawn by the computer at the beginning of the game.

Players know that they are player X or Y. But only Y knows that she/he is BLUE or YELLOW. Y’s status is unknown to X player. In each round, 20 players are divided into 10 pairs. The table below shows a possible allocation of 10 pairs:

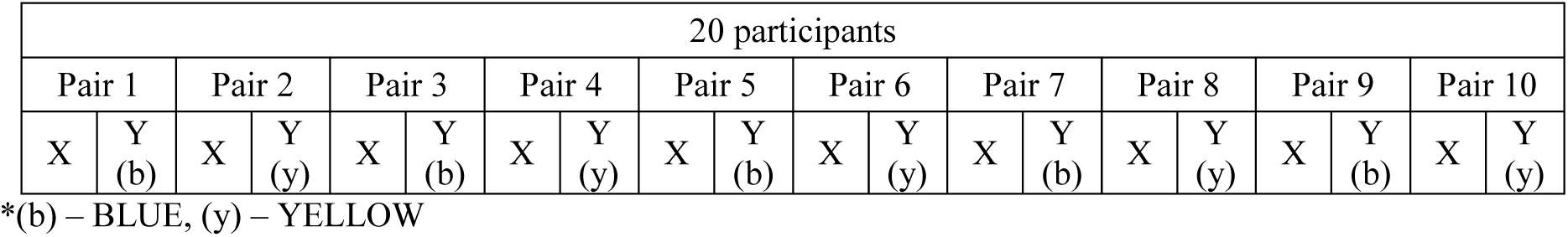

The game is the following: X owns a resource, and Y wants to obtain that resource. If Y succeeds to request the resource from the player X - whether Y is the BLUE or the YELLOW - it means HUF 1200 to them. If the player Y fails to get the money, she/he gets HUF 0. The possible payoffs of Y are shown in the following table:

#### Y’s payoffs along with different decisions

**Y's payoffs along with different decisions:**
|  | X's decision |  |
| --- | --- | --- |
| Y state: | Give the resource | Do not give the resource |
| BLUE | HUF 1200 | HUF 0 |
| YELLOW | HUF 1200 | HUF 0 |

X does not know the state of Y. However, X wins only in those cases if she/he gives the resource to Y who is in the BLUE state or does NOT give the resources to those Ys who are in the YELLOW state. The possible payoffs of X are shown in the following table. If player X gives the resource to player Y in the BLUE state, X wins HUF 1200, but if X gives the resource to player Y in the YELLOW state, X does not get anything. Player X also earns HUF 1200 if X does NOT give the resource to the YELLOW players, and conversely, if player X does not give the resource to Ys in the BLUE state X does not win anything.

#### X’s payoffs along with different decisions

**X's payoffs along with different decisions:**
|  | X's decision |  |
| --- | --- | --- |
| Y state: | Give the resource | Do not give the resource |
| BLUE | HUF 1200 | HUF 0 |
| YELLOW | HUF 0 | HUF 1200 |

Y player can use two signals to request the resource from X:

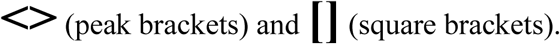

These marks may reduce, increase, or make no changes on the payoff of player Y in the game. The way it affects the payoffs will appear next to the sign during the game.

##### Example 1

Y can select from the following signs: (1) using the <> sign reduces Y’s payoff by HUF 300; (2) using the [] sign increases Y’s payoff by HUF 300. Y decided to use the <> sign and X gave him the resource. In this case, the payoff of Y will be 1200 - 300 = 900.

##### Example 2

Y can select from the following signs: (1) using the <> sign has no effect on Y’s payoff; (2) using the [] sign reduces Y’s payoff by HUF 500. Y decided to use the <> sign and X gave him the resource. In this case, the payoff of Y will be 1200 - 0 = 1200.

##### Example 3

Y can select from the following signs: (1) using the <> sign reduces Y’s payoff by HUF 300; (2) using the [] sign increases Y’s payoff by HUF 300. Y decided to use the [] sign and X gave him the resource. In this case, the payoff of Y will be: 1200 + 300 = 1500.

##### Example 4

Y can select from the following signs: (1) using the <> sign reduces Y’s payoff by HUF 300; (2) using the [] sign increases Y’s payoff by HUF 300. Y decided to use the <> sign and X did NOT give him the resource. In this case, the payoff of Y will be 0 - 300 = −300.

Each round is important since, at the end of the game, participants will be paid on the basis of two randomly selected rounds where they played role X and Y. If a participant’s payoff is negative, then we rounded the amount up to HUF 0. In addition to this calculated payoff, participants receive a show-up fee of HUF 1000.

The time available for your choice will display in the top right corner of the screen.

Thank you for your participation!

Have fun and good luck!

### FURTHER INSTRUCTION I

In the next game, Player Y can use one of the following signals depending on their color to get the resource from player X. Under the signals you can see how each signal can change your earnings.

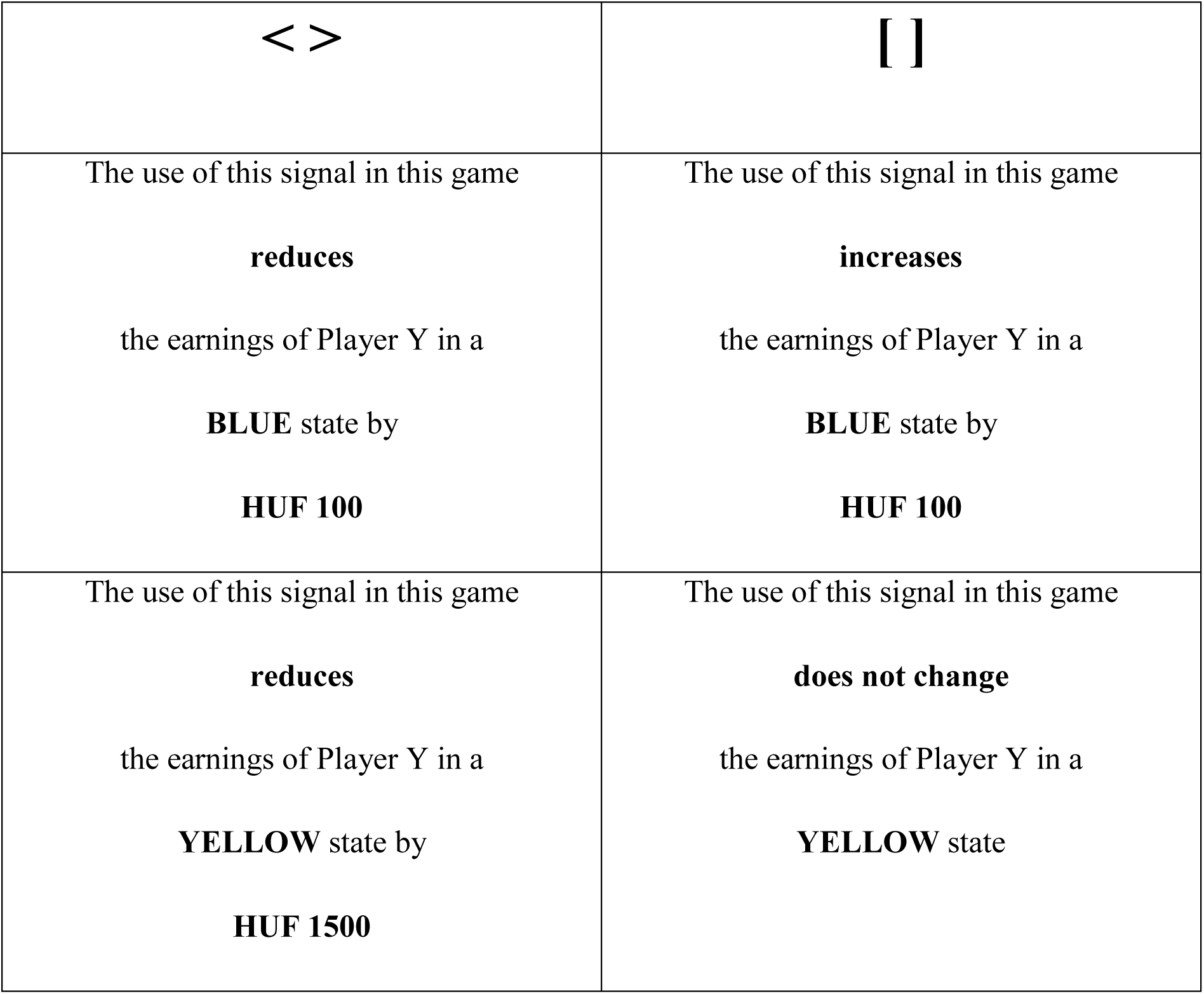

